# Development of a Method Combining Peptidiscs and Proteomics to Identify, Stabilize and Purify a Detergent-Sensitive Membrane Protein Assembly

**DOI:** 10.1101/2022.03.03.482899

**Authors:** John William Young, Irvinder Singh Wason, Zhiyu Zhao, Sunyoung Kim, Hiroyuki Aoki, Sadhna Phanse, David G Rattray, Leonard J. Foster, Mohan Babu, Franck Duong van Hoa

**Author notes:** These authors contributed equally to this work (John William Young, Irvinder Singh Wason).

## Abstract

The peptidisc membrane mimetic enables global reconstitution of the bacterial membrane proteome into water-soluble detergent-free particles, termed peptidisc libraries. We present here a method that combines peptidisc libraries and chromosomal-level gene tagging technology with affinity purification and mass spectrometry (AP/MS) to stabilize and identify fragile membrane protein complexes that exist at native expression levels. This method circumvents common artifacts caused by bait protein overproduction and protein complex dissociation due to lengthy exposure to detergents during protein isolation. Using the *E. coli* Sec system as a case study, we identify an expanded version of the translocon, termed the HMD complex, consisting of 9 different integral membrane subunits. This complex is stable in peptidiscs but dissociates in detergent. Guided by this native-level proteomic information, we design and validate a procedure that enables purification of the HMD complex with minimal protein dissociation. These results highlight the utility of peptidiscs and AP/MS to discover and stabilize fragile membrane protein assemblies.

## INTRODUCTION

Early efforts to characterize membrane protein interaction networks – or the “membrane protein interactome” – have mostly relied on the over-production of bait proteins followed by membrane solubilization with detergents (*1–5*). After isolation of the bait, potential interactors are identified by mass spectrometry or immunoblotting (*1–3, 5*). Although straightforward, there are major drawbacks to this method. First, detergent micelles, which are necessary to maintain membrane protein solubility, tend to dissociate fragile multisubunit complexes, causing protein-protein associations to be lost (*1–4, 6–8*). Detergents must also be carefully removed before mass spectrometry analysis, which tends to decrease the efficiency of protein identification. Second, plasmid-based over-expression of the bait protein often perturbs cell envelope biogenesis and potentially alters the stoichiometry and specificity of protein interactions, leading to adverse effects on cell physiology and complex identification (*4, 9, 10*).

Several chromosomal-level tagging approaches have been developed recently to bypass the need for bait protein overproduction (*1, 11, 12*). Work from our group and others has shown the utility of chromosomal tagging approaches to map protein interaction networks in bacteria, yeast, and mammalian cells (*1, 2, 11–13*). To circumvent the adverse effects of detergents, several membrane mimetics have been developed to isolate membrane proteins in a water-soluble state (*4, 14–16*). Results from our group have shown that the peptidisc enables reconstitution of a significant fraction of the membrane proteome in a form suitable to mass spectrometry analysis (Figure 1) (*4, 16*).

**Figure 1:**
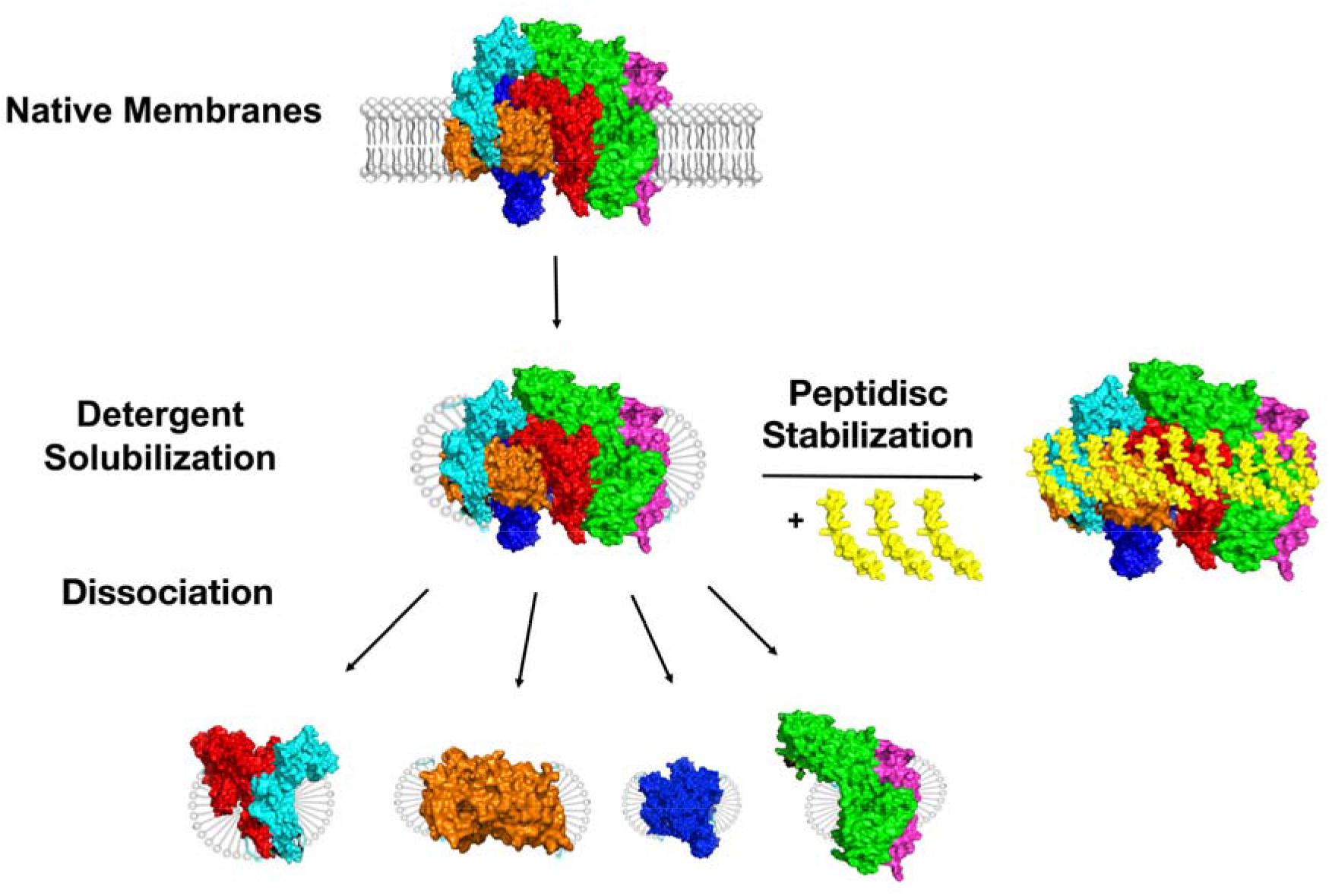
Stabilization of multi-subunit membrane protein complexes using the peptidisc. A multi-subunit membrane protein complex is initially extracted from the lipid bilayer with detergents. However, the complex is prone to dissociation into sub-complexes following prolonged detergent exposure. Peptidisc reconstitution enables stabilization of the assembly for downstream purification.

In this report, we combine these two recent developments to identify and stabilize membrane protein interactions at native expression levels, and we use this information to guide the development of a protocol to purify a membrane protein complex of interest with minimal interference from detergent. We start with an *E. coli* strain that is modified at the chromosomal-level with a sequential peptide affinity (SPA) tag inserted at a specific gene position, thereby creating a C-terminal fusion with the protein of interest. (*1, 10*). After reconstitution of the membrane proteome into peptidiscs, the bait and its co-purifying interactors are identified by AP/MS analysis. To solidify our initial findings, and to better characterize the interactome of our protein of interest, we then repeat this peptidisc-AP/MS workflow using SPA-tagged strains for the most prominent interactors identified in our initial experiment. To differentiate between enriched interactors and non-specific protein contaminants, we performed this workflow in parallel using an *E. coli* strain with no SPA tag.

As a case study, we apply this Peptidisc-AP/MS workflow toward characterizing the membrane interactome of the highly conserved bacterial translocon - also known as the SecYEG complex - which forms a protein-conducting channel across the inner membrane in gram-negative bacteria (*17, 18*). We selected the translocon as an experimental testbed because it is known to associate dynamically with a wide variety of integral membrane protein subunits. These include the membrane-integrated SecDFyajC complex and the membrane insertase YidC, as well as the membrane-tethered periplasmic chaperones YfgM and PpiD (*3, 4, 19–23*). Previous biochemical studies have isolated a multi-subunit complex - termed the bacterial “holo-translocon” (HTL) - consisting of SecYEG, SecDFyajC, and YidC (*22, 24–26*). However, the HTL purification is technically challenging, requiring multiple different tags and chromatography steps, as well as additional specific phospholipids to minimize the dissociating effects of detergent (*22, 24, 27*).

Using Peptidisc-AP/MS workflow described above, we identify an expanded version of the holo-translocon in the native membrane, which we term the “HMD” complex, consisting of SecYEG, the SecDFyajC, and YidC subcomplexes, plus the periplasmic chaperones YfgM and PpiD. Having shown that this assembly can be isolated in peptidiscs, we use this information to develop a protocol that enables overproduction and biochemical purification with minimal protein dissociation.

## MATERIALS AND METHODS

### Reagents

Tryptone, yeast extract, Na_2_HPO_4_, KH_2_PO_4_, NaCl, and Tris-base were obtained from Bioshop Canada. Ampicillin, kanamycin, and arabinose were purchased from GoldBio. n-dodecyl-β-d-maltoside (DDM) was purchased from Anatrace. ANTI-FLAG M2 affinity gel was purchased from Sigma. Peptidiscs (purity >80%) were obtained from Peptidisc Biotech Canada. All other chemicals were obtained from Fisher Scientific Canada.

### Plasmids and strains

*E. coli* SPA (Sequential Peptide Affinity)-tagged strains were from the Babu laboratory collection (*1*). The strains used in this study were as follows: DY330 (non-tagged parental control strain), b3300 (SecY-SPA), b0408 (SecD-SPA), b0441 (PpiD-SPA), and b2513 (YfgM-SPA). Plasmids pBad22-*hisEYG*, pBad33-*DFyajC*, pBad22-*hisYidC*, and pBad33-*YfgM_His_-PpiD* have been previously described (*4, 5, 28–30*). All plasmids generated in this study were cloned using the PIPE method (*31*). The genes for SecDF were amplified from pBad33-*DFyajC* and inserted into pBad22-*hisEYG*, generating the plasmid pBad22-*hisEYGDF*. To express the bacterial holo-translocon (HTL; a complex of SecYEG with SecDFyajC and YidC (*22*)), the gene for YidC without its affinity tag was amplified from pBad22-*hisYidC* and inserted into pBad22-his*EYGDF* to form the plasmid pBad22-*HTL*. The His-tag was deleted from pBad33-*YfgM_His_-PpiD* to generate pBad33-*YfgM-PpiD*. All constructs were verified by DNA sequencing (Genewiz).

### Growth of SPA-tagged strains and membrane preparation

All strains except the DY330 control strain were revived on Luria Bertani (LB)-agar plates supplemented with 25 μg/mL kanamycin. Strain DY330 was revived on a LB-agar plate without antibiotics. For each strain, a single colony was used to inoculate a 10 mL overnight culture in LB media (plus antibiotic, where appropriate). Overnight cultures were grown overnight at 37 °C with shaking. The following day, overnight cultures were diluted 1/100 into 1L fresh LB media and grown for a further 6 hours until OD_600_ ~1. Cells were harvested by centrifugation (6,000 x g, 10 minutes) and resuspended in TSG (50 mM Tris HCl pH 8; 50 mM NaCl; 10% glycerol) buffer containing 1 mM PMSF before being Dounce homogenized and lysed on a Microfluidizer (12,000 psi, three passes). Lysates were centrifuged at 6,000 x g for 10 minutes to remove unbroken cells. The membrane fraction was collected by ultracentrifugation (100,000 x g, 30 minutes) in a Ti70 rotor. After centrifugation, the supernatant was discarded, and the membrane pellet was resuspended in TSG buffer.

### Preparation of peptidisc libraries

Membranes (~1-2 mg) were solubilized with 0.5% DDM in 1 mL volume on ice for 15 minutes. Insoluble material was removed by ultracentrifugation (100,000 x *g*, 15 minutes). The detergent-solubilized material was immediately mixed with a 4:1 excess of NSP_R_ peptidisc peptide in a 15 mL 100 kDa cutoff Amicon concentrator. To form a peptidisc library, the solubilized membrane-peptide mixture was diluted to 10 mL in TSG buffer. The mixture was concentrated to ~1 mL by centrifugation (3,000 x g, 10 minutes) at 4°C before being diluted back to 10 mL and re-concentrated to ~1 mL.

### FLAG pulldowns in peptidisc

About 1 mg peptidisc libraries were loaded onto 50 μL ANTI-FLAG M2 affinity gel and incubated at 4°C overnight with gentle shaking. The following day, the flow-through was collected, and the resin was washed with 1 mL TSG buffer. Five additional 1 mL washes were performed to minimize non-specific binding to the resin. Bound proteins were eluted in 100 μL 100 mM Glycine HCl pH 3.5. The sample pH was adjusted by adding 10 μL 1M Tris HCl pH 8.0. Samples were then digested and STAGE tipped before being analyzed by LC-MS/MS exactly as previously described (*1*).

### FLAG pulldowns in detergent

Membranes were solubilized as described above. About 1 mg of detergent-solubilized material was loaded onto 50 μL ANTI-FLAG M2 affinity gel and incubated at 4°C overnight with gentle shaking. The next morning, the flow-through was collected, and the resin was washed with 1 mL TSG buffer + 0.02% DDM. Five additional 1 mL washes were performed to minimize non-specific binding to the resin. Bound proteins were eluted in 100 μL 100 mM Glycine HCl pH 3.5. Sample pH was adjusted by adding 10 μL 1M Tris HCl pH 8. The samples were precipitated with ice-cold acetone to remove detergent (overnight incubation, 4°C). Precipitated proteins were pelleted by centrifugation (10,000 g, 10 minutes), and the supernatant was removed using a vacuum aspirator. Protein pellets were dried at 42°C for 10 minutes. Samples were then digested and STAGE tipped before being analyzed by LC-MS/MS exactly as previously described (*1*).

### Analysis of mass spectrometry data

Analysis was performed using MaxQuant version 1.6.17.0 (*32, 33*). The search was performed against a database comprised of the protein sequences from the source organism (*E. coli* K12) plus common contaminants using the following parameters: peptide mass accuracy <5 ppm; fragment mass accuracy 0.006 Da; trypsin enzyme specificity; fixed modifications, carbamidomethyl; variable modifications, methionine oxidation, deamidated N, Q, and N-acetyl peptides. Proteins were quantified from 1 peptide identification. Only those peptides exceeding the individually calculated 99% confidence limit (as opposed to the average limit for the whole experiment) were considered as accurately identified.

### Capturing labile interactors of the SecYEG complex in peptidiscs

To over-produce the SecYEG and HTL complexes, plasmids pBad22-*EYG* and pBad22-*HTL* were transformed into chemically competent BL21(DE3) cells. To facilitate over-production of the whole “HMD” complex, the plasmid pBad33-*YfgM PpiD* (containing no affinity tag) was transformed into BL21(DE3) competent cells containing the plasmid pBad22-*HTL*. Protein expression and membrane isolation were performed as previously described (*28, 34, 35*). Briefly, protein expression was induced for 3 hr at 37°C after induction with 0.2% Arabinose at an OD of 0.4–0.7 in 1L LB medium supplemented with appropriate antibiotics. Cell lysis and membrane isolation were as described above. To enrich the inner membrane further, the membrane fraction was layered onto a 2-step 20%-50% sucrose gradient in SW41 tubes and re-centrifuged at 200,000 *g* for 2 hours. Inner Membrane Vesicles (IMVs) were recovered as a distinct brown band near the middle of the gradient and diluted 4-fold in TSG before being pelleted by ultracentrifugation (100,000 x *g*, 15 minutes). To verify successful over-production of all protein subunits, a 5-μg aliquot of each membrane preparation was analyzed by 15% SDS-PAGE followed by Coomassie Blue staining.

For purifications in detergent conditions, IMVs were solubilized with 0.5% DDM in 1 mL volume on ice for 15 minutes. Insoluble material was removed by ultracentrifugation (100,000 x *g*, 15 minutes). A 1 mL aliquot of the detergent extract was incubated with 50 μL Ni-NTA resin for 30 minutes. The resin was washed three times with 10 column volumes (CVs) of TSG buffer + 0.02% DDM, then eluted in TSG buffer + 0.02% DDM + 300 mM Imidazole. Eluted proteins were analyzed by 15% SDS PAGE and visualized by Coomassie Blue staining. For reconstitution into peptidiscs, a freshly solubilized detergent extract was mixed with a 4:1 excess of peptidisc peptide in a 15 mL 100 kDa cutoff Amicon concentrator and reconstituted into peptidisc libraries as previously described (*4*). The resultant library was injected onto a Superose 6 10/300 column (GE Healthcare) equilibrated in TSG buffer on an AKTA FPLC. Peak fractions were analyzed by SDS-PAGE to verify the presence of all protein subunits. Peak fractions were then pooled and incubated with Ni-NTA resin for 30 minutes at 4°C. The resin was washed three times with 10 CVs of TSG buffer, then eluted in TSG buffer + 300 mM Imidazole. Eluted proteins were analyzed by 15% SDS PAGE and visualized by Coomassie Blue staining.

## RESULTS

### An AP/MS approach to identify the interactome of the Sec translocon

To benchmark our Peptidisc-AP/MS workflow and identify the SecYEG complex’s primary interactors in native membranes, we employed a strain with the sequential peptide affinity (SPA) tag inserted in the *secY* gene (strain termed SecY-SPA). This strain is perfectly viable, indicating that the SecY protein modified with a C-terminal SPA tag is functional (*1*). Following cell lysis, the total envelope proteome containing inner and outer membrane proteins was briefly solubilized with n-dodecyl-β-d-maltoside (DDM). After removing aggregates by ultracentrifugation, a 4:1 mass excess of peptidisc peptide was added, and the detergent micelles were rapidly removed via successive dilution and filtration steps as described in the Materials and Methods. The resulting peptidisc library was then affinity purified over anti-FLAG resin, and the eluate was digested and STAGE tipped before being analyzed in triplicate by LC-MS/MS. A total of 240 proteins were identified in all three replicates (Supplemental File 1). As a graphic representation of these data, the protein list was ranked in alphabetical order, and the intensity value of each protein was plotted as a function of its rank in the list (Figure 2A). To enable identification of the protein contaminants inherent to this type of analysis (e.g., non-specific interactors of the anti-FLAG resin), the peptidisc AP/MS workflow was done in parallel using a library prepared from the parental wild-type strain DY330 (*1*). In this control, a total of 266 proteins were identified in all three replicates. The protein list was ranked and plotted as described above (Figure 2B).

**Figure 2:**
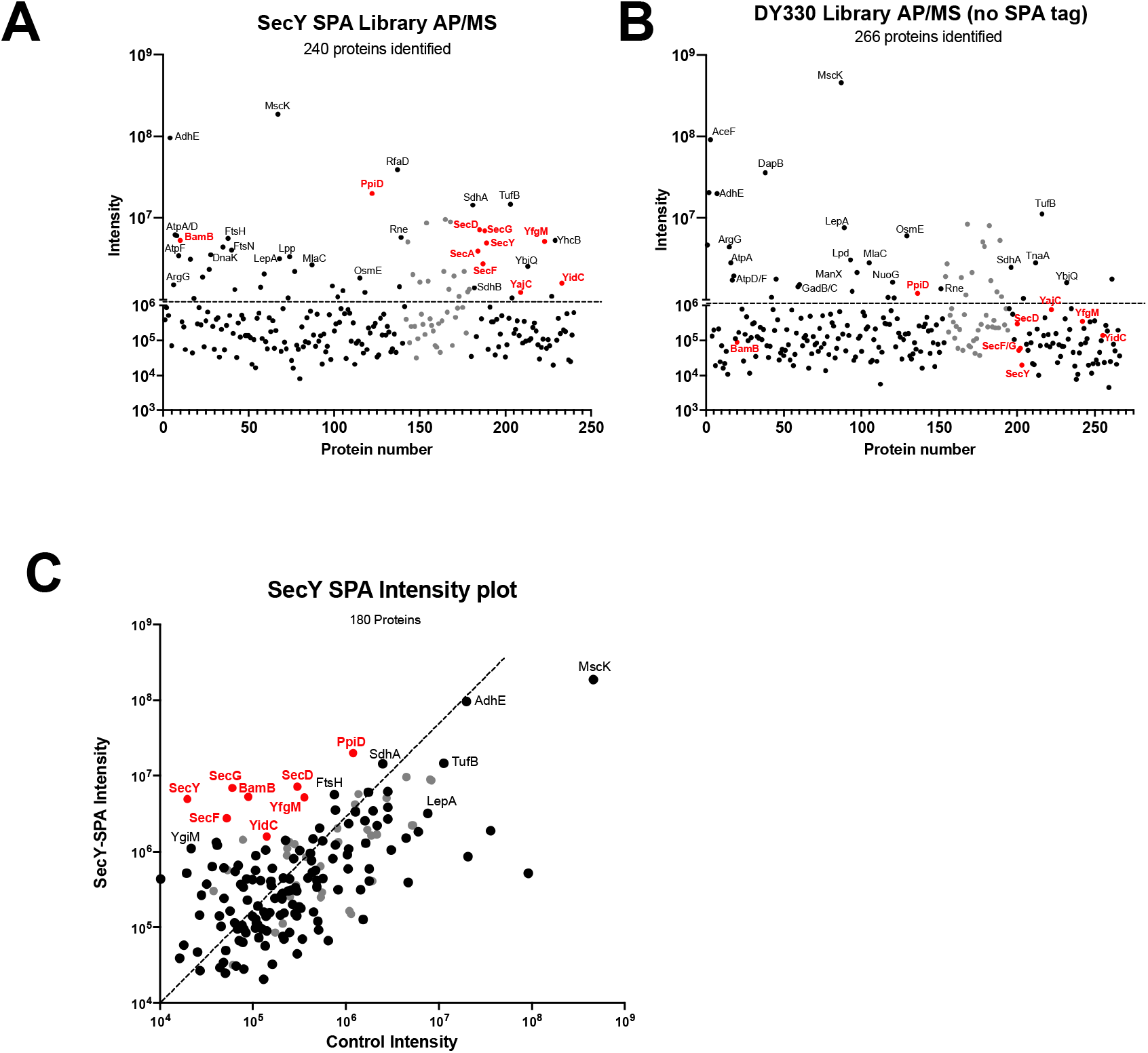
Capture and identification of SPA-tagged SecY using FLAG antibodies. **A.** Peptidisc libraries prepared from an *E. coli* strain modified with a chromosomally SPA-tagged SecY protein were purified over ANTI-FLAG M2 Agarose beads. LC-MS/MS identified eluted proteins. The protein list was ranked alphabetically to visualize the data, and protein intensities were plotted against protein rank in the list. Proteins of interest are highlighted in red; ribosomal subunits are in grey; all other identified proteins are shown in black. **B.** To identify non-specific protein contaminants, a peptidisc library prepared from the parental strain (no SPA tag) was processed in parallel. Data were analyzed and plotted as in (A). **C.** To give an idea of protein enrichment, the intensity of each protein present in the SecY sample was plotted against its intensity in the control sample. Colour coding was as described in (A).

Comparing the two plots side-by-side, it appears that the SecYEG translocon ancillary membrane subunits PpiD, YfgM, SecD, SecF, SecG, and YidC, as well as SecY itself, are present at far higher intensities in our SecY-SPA pulldown compared to the control. The BamB subunit of the Bam complex is also enriched (red points, Figures 2A and 2B). We note that multiple additional proteins are present at comparable intensities between the SecY-SPA and control pulldowns - including MscK, TufB, SdhA, and multiple ATP synthase subunits (black points, Figures 2A and 2B). Numerous cytosolic ribosomal subunits are also present in both pulldowns (grey points, Figure 2A and 2B). The detection of numerous protein contaminants is inherent to AP/MS experiments and proteomic analysis in general (*1, 3, 4*).

As an alternative representation of our data, we next plotted the intensities of each protein present in both the SecY-SPA and control pulldowns on the same graph (Figure 2C). Presenting the data in this manner makes it easier to identify specifically enriched proteins versus non-specific contaminants. The result shows that 180 proteins are present in both pulldowns and, consistent with our earlier analysis, the known Sec translocon subunits PpiD, YfgM, SecD, SecF, SecY, SecG, and YidC are clearly enriched (red points). Abundant contaminants such as MscK, TufB, SdhA, and multiple ribosomal subunits are present at roughly equal intensities in both pulldowns.

As a cross-validation assay, we applied our peptidisc AP/MS workflow to three additional strains, encoding for SPA-tagged SecD, PpiD, YfgM. Data were analyzed and visualized as described above (Figure 3A-C). We note that the extent of non-specific protein contamination appears somewhat higher in the SecD and YfgM FLAG pulldown experiments compared to the PpiD FLAG pulldown. However, despite the presence of these protein contaminants, we find these latest results are confirmatory of the findings in our earlier SecY peptidisc AP/MS and suggest a network of interactions between the core Sec translocon, SecD, SecF, YidC, YfgM, and PpiD.

**Figure 3:**
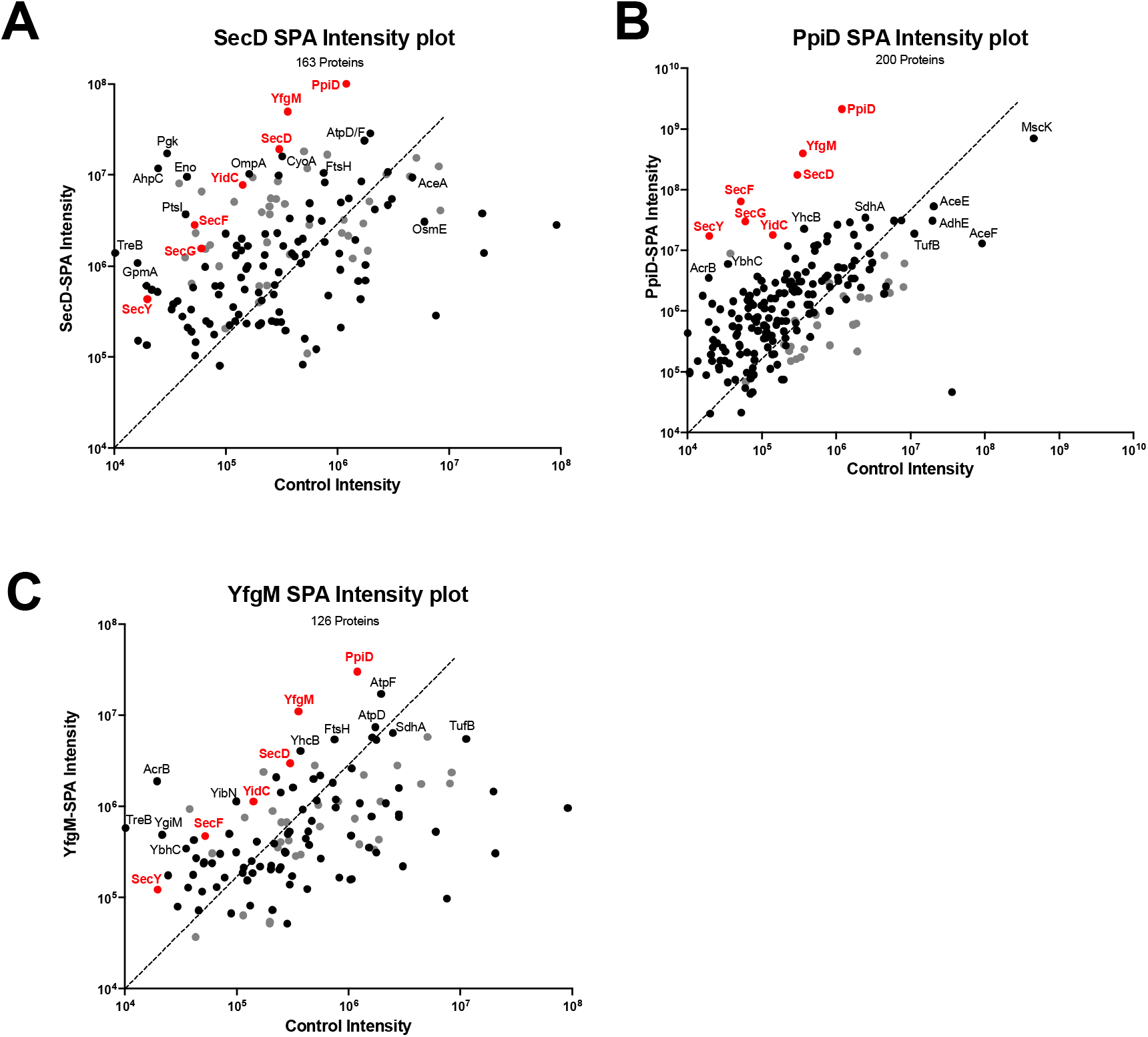
Capture and identification of Sec translocon ancillary subunits by FLAG AP/MS. **A.** Capture and identification of SPA-tagged SecD using FLAG antibodies. Peptidisc libraries prepared from an *E. coli* strain modified with a chromosomally SPA-tagged SecD protein were purified over ANTI-FLAG M2 Agarose beads. Data were analyzed and visualized as in Figure 4C. **B.** As in (A), but for SPA-tagged PpiD. **C.** As in (A) but for SPA-tagged YfgM.

To compare the peptidisc AP/MS workflow against a classical detergent-based AP/MS approach, we performed AP/MS using a detergent extract prepared from the strain SPA-tagged for PpiD as described in the Materials and Methods (Figure 4A). PpiD was selected as the bait because its other Sec interactors are well detected in our peptidisc AP/MS experiments above. The experiment was done in parallel using a detergent extract from the parental DY330 strain as a control (Figure 4B). As expected, the results with the PpiD detergent extract show that the protein YfgM is enriched along with PpiD (Figure 4A and 4C) (*3*). However, many interactors of PpiD identified in our peptidisc AP/MS experiment above - including SecD, SecF, YidC, and SecYEG - are either poorly detected in detergent (i.e., with low intensity) or are undetected. Likely, these interactions are mostly dissociated upon prolonged exposure to detergents, particularly during the washing steps, which are necessary to reduce the abundance of non-specific protein contaminants. We did not observe this protein dissociation in our peptidisc experiments, likely because the reconstitution step to stabilize fragile complexes is performed before affinity purification and because washing is performed without detergents.

**Figure 4:**
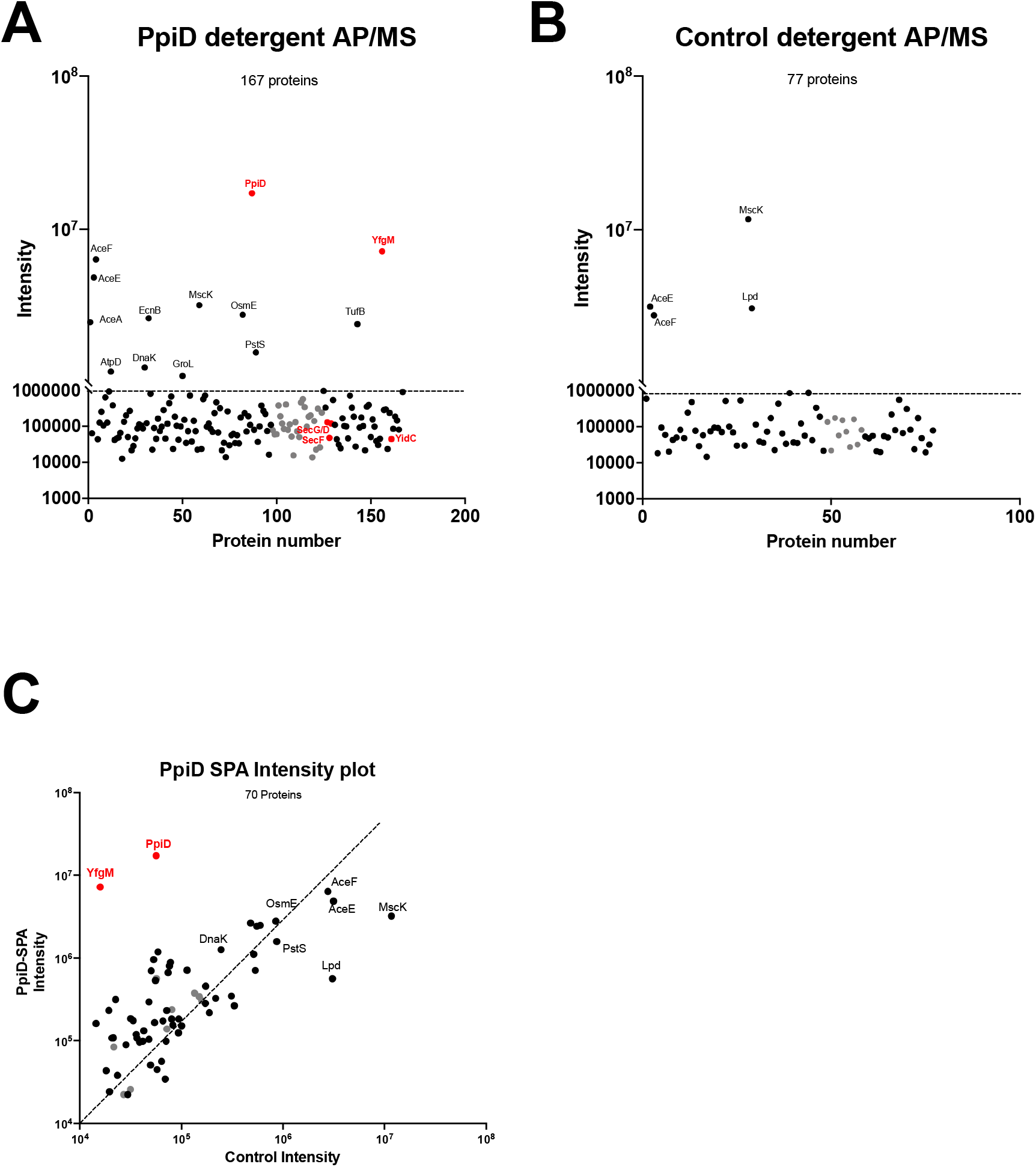
Capture and identification of SPA-tagged PpiD in DDM using FLAG antibodies. **A.** A detergent extract prepared from an *E. coli* strain modified with a chromosomally SPA-tagged PpiD protein was purified over ANTI-FLAG M2 Agarose beads. Data were analyzed and visualized as described in Figure 2A. **B.** To identify non-specific protein contaminants, a detergent extract prepared from the parental strain (no SPA tag) was processed in parallel. Data were analyzed and plotted as in (A). **C.** To give an idea of protein enrichment, the intensity of each protein present in the PpiD sample was plotted against its intensity in the control sample. Colour coding was as described in (A).

### Biochemical isolation of the HMD complex in peptidiscs

Altogether, our proteomic data reveals interactions between SecYEG, SecDF, YidC, YfgM, and PpiD in native *E. coli* membranes. This network of interactions - identified here across multiple peptidisc AP/MS experiments using multiple different baits – likely represents a super-complex containing all subunits of the previously characterized bacterial holo-translocon (SecYEG, SecDF, and YidC), plus the membrane-tethered chaperones YfgM and PpiD. Thus, we are naming this complex the “HMD” (<u>h</u>olo-<u>t</u>rans<u>l</u>ocon plus Yfg<u>M</u> and Ppi<u>D</u>) complex. Guided by this information, we next designed a protocol to enable its biochemical purification.

We cloned the holo-translocon (HTL) SecYEG-SecDF-YidC subunits into a bacterial expression vector. The protein SecE bears an N-terminal His-tag to facilitate affinity purification of the complex. For over-production of the entire HMD complex, non-tagged YfgM and PpiD were cloned into a second plasmid with different antibiotic resistance.

To verify successful over-production of all protein subunits, we expressed the HTL and HMD complexes in *E. coli* BL21(DE3) and isolated the inner membranes. In addition, the core translocon SecYEG was expressed side-by-side as a control, and to visualize protein content, the membranes were analyzed by 15% SDS-PAGE and Coomassie blue staining (Figure 5A). Encouragingly, we observe that all protein subunits of the HMD complex are produced in roughly similar amounts.

**Figure 5:**
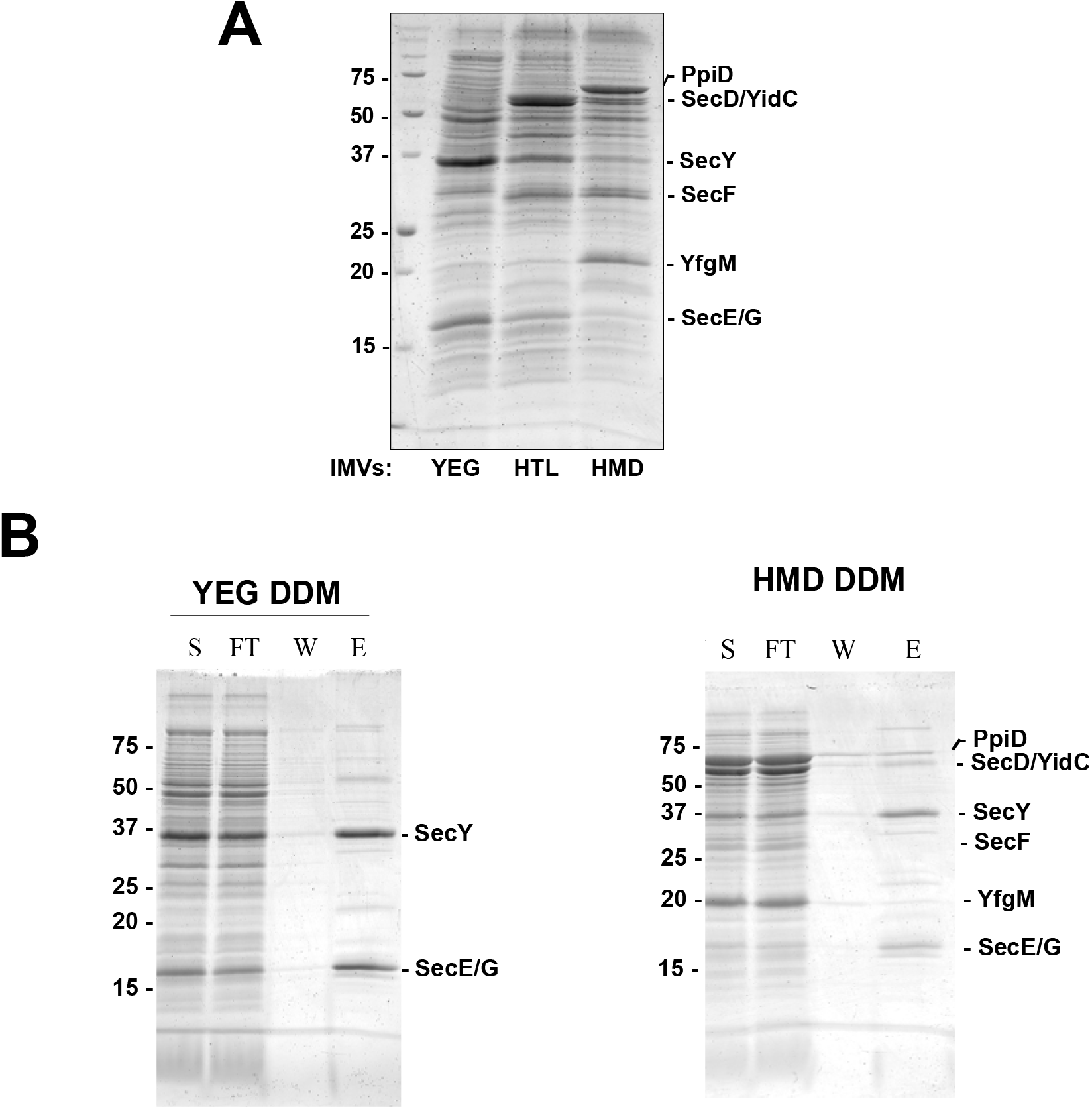
Detergent sensitivity of the “HMD” complex. **A.** SDS-PAGE analysis of *E. coli* inner membranes containing the over-expressed SecYEG, HTL, and HMD complexes. Migration of each over-expressed protein subunit is indicated on the right side of the gel. **B.** Membranes containing SecYEG and HMD from (A) were solubilized in DDM and purified over Ni-NTA affinity resin. Aliquots of each purification were analyzed by SDS-PAGE and Coomassie staining (S = untreated start material; FT = unbound flowthrough; W = wash; E = elution).

Membranes bearing the SecYEG and HMD complexes were solubilized briefly in DDM and affinity-purified in detergent. Aliquots from each purification step were analyzed by SDS-PAGE and Coomassie blue staining (Figure 5B). Consistent with our initial proteomic observations, the HMD complex appears largely dissociated, with the PpiD, YfgM, SecDF, and YidC subunits primarily lost in the detergent washing steps. On the other hand, the core translocon SecYEG appears stable during purification in DDM, as expected.

Next, we reconstituted a fresh aliquot of the unpurified HMD detergent extract into peptidiscs. To verify the solubility of our preparation, we first fractionated the sample by size exclusion chromatography (SEC) in detergent-free conditions. Visual examination of the SEC chromatogram reveals two peaks - the first corresponds to the reconstituted peptidisc library, while the second corresponds to excess peptidisc peptides (Figure 6A). The peptidisc library clearly elutes after the column’s void volume, indicating that the library is watersoluble and free of aggregates. Fractions under the central peak were analyzed by SDS-PAGE (Figure 6B). As expected, all subunits of the HMD complex are present in roughly stoichiometric amounts. These peaks fractions were pooled, and the HMD complex was isolated by Nickel affinity purification. Aliquots from each purification step were analyzed by SDS-PAGE (Figure 6C). Consistent with our earlier proteomic observations, the PpiD, YfgM, SecDF, and YidC subunits co-purify along with SecYEG.

**Figure 6:**
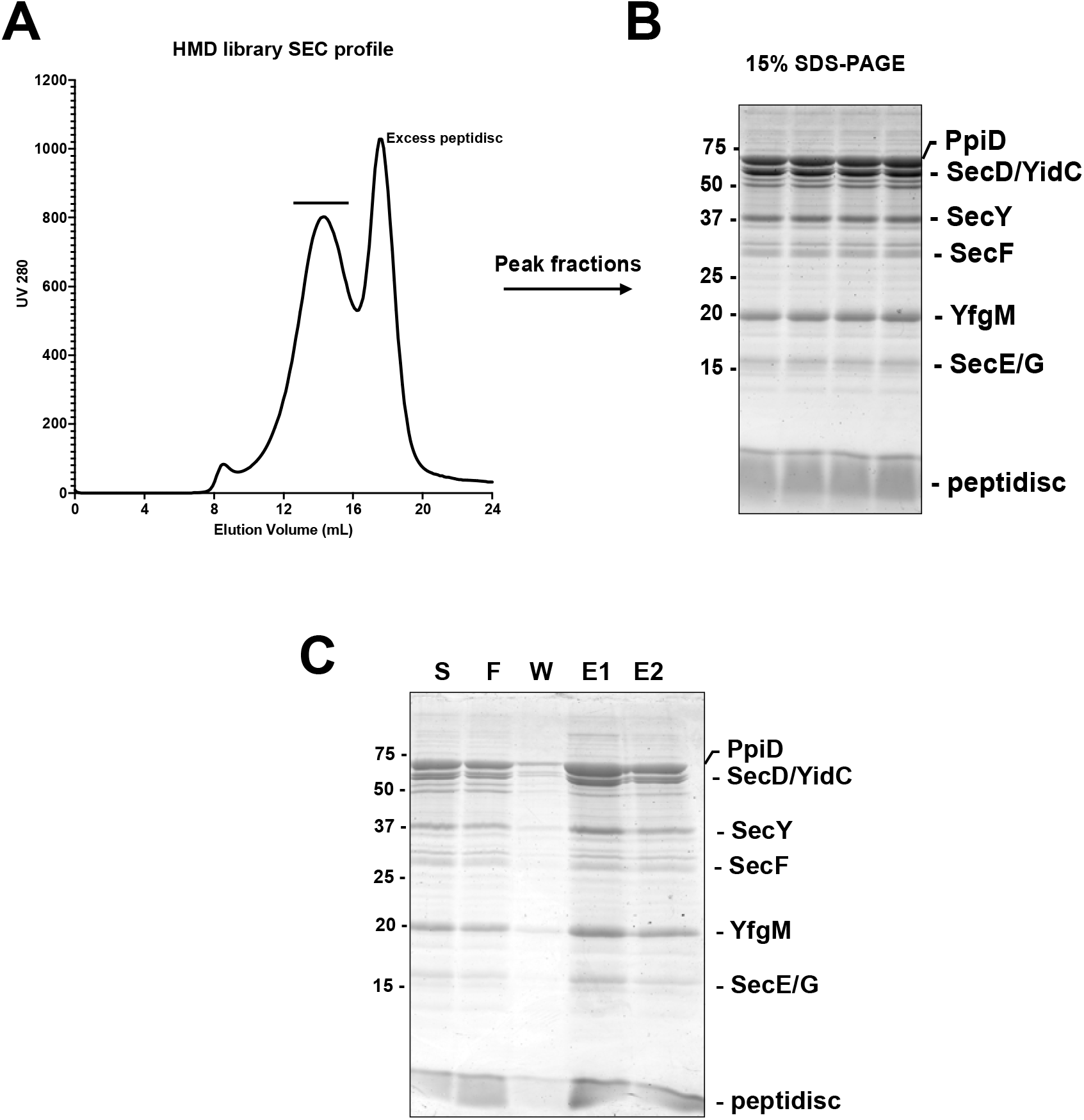
Purification of the “HMD” complex. **A.** A peptidisc library containing the HMD complex was injected on a Superose 6 10/300 column equilibrated in a detergent-free buffer. Protein abundance was monitored by UV (280 nm) and plotted as a function of elution volume (mL). Large aggregates are expected to elute at 8 mL - the void volume of the column. **B.** Fractions under the central peak (indicated by a black bar in (A) were analyzed by SDS-PAGE and Coomassie staining. Migration of over-expressed protein subunits is indicated on the right-hand side of the gel. **C.** Peak fractions from (B) were pooled, and the HMD complex was isolated by affinity pulldown.

## DISCUSSION

In this study, we develop a method to detect multi-subunit membrane protein complexes which are naturally present in the cell membrane, yet difficult to identify because of their low abundance and facile dissociation when extracted with detergent. Specifically, we combine the peptidisc membrane mimetic with a series of *E. coli* strains that are modified with a sequential peptide affinity (SPA) tag at the chromosomal-level. This approach maintains expression of the target protein at near-endogenous levels, which minimizes artifacts due to protein overproduction such as misfolding and mis-localization (*1, 10*). This approach also decreases protein complex dissociation because their capture peptidiscs allows to minimize exposure to detergents. After stabilization of the membrane complex in peptidiscs, the SPA-tagged bait proteins and their interactors are isolated by affinity chromatography and identified by mass spectrometry. To facilitate identification of background contaminants versus enriched interactors, the pulldown experiment is repeated in parallel using a parental *E. coli* strain with no SPA tag.

Using the well-characterized bacterial Sec translocon as an experimental model, we perform a series of AP/MS experiments on multiple known Sec ancillary subunits. This work led us to identify an expanded version of the holo-translocon complex consisting of SecYEG, SecDFyajC, YidC, plus the two membrane chaperones YfgM and PpiD. We named it the “HMD complex” for <u>h</u>olo-<u>t</u>rans<u>l</u>ocon plus Yfg<u>M</u> and Ppi<u>D</u>. Guided by these proteomic observations, we develop a purification protocol to isolate biochemical amounts of the HMD complex. We start by overproducing the HMD complex with a single affinity tag using *E. coli* expression vectors. We note that the over-produced HMD complex is largely dissociated when attempting purification in detergent conditions. Even mild detergents such as DDM are known to exhibit dissociative and delipidating effects on membrane proteins, particularly during the washing steps that are intrinsic to affinity purification (*4, 36, 37*). Therefore, to minimize these detergent effects, we reconstitute the HMD complex into peptidiscs before attempting any purification steps. When analyzed by size-exclusion chromatography (SEC) in detergent-free conditions, we observe that our peptidisc preparation is water-soluble and free of aggregates and that all subunits of the HMD complex co-elute together from the column, indicating that the complex is stabilized in peptidiscs. After pooling the peak fractions from our initial SEC experiment, we can then isolate the HMD complex by affinity purification.

Although this study does not reveal novel protein interactors of the Sec translocon, this work provides the first evidence that the “HMD” complex is an entity present in the native membrane. Other studies have determined that the copy number of SecYEG and interacting partners differs considerably within the cell (*21, 38*). Given these different expression levels, the “HMD” complex identified here is likely to be a highly dynamic assembly in native membranes, rapidly assembling and disassembling into various sub-complexes. This dynamic association would explain the fragility of the complex. Considering this, we do not exclude the possibility that our purified preparation contains a mixture of both complete “HMD” complex and varied sub-complexes. Details regarding the stoichiometry, dynamics, and structure of the HMD complex will need to be presented in future experiments. The identity of any annular lipids that may modulate the HMD complex’s activity may also be a promising avenue for future investigation.

This current work represents the results of a pilot study using a series of SPA-tagged *E. coli* strains. These strains are only a tiny subset of a more extensive strain library available in our laboratory, in which hundreds of *E. coli* membrane protein open-reading frames (ORFs) have been systematically SPA-tagged. A similar public library is also available to characterize the *S. cerevisiae* membrane proteome (*1, 2*). Other recent work reports the high throughput construction of new strains and cell lines to map protein interaction networks in mitochondria and mouse brain tissue using AP/MS (*13, 39, 40*). Given the widespread availability of geneediting technology, we anticipate that our Peptidisc-AP/MS approach will be easily expandable to precisely characterize membrane protein interactomes across multiple different organisms and tissue types.

## ACKNOWLEDGEMENTS

This work was supported the Canadian Institutes of Health Research to the FDVH’s group. Platform funding for proteomics research in L.J.F.’s group is supported by Genome Canada and Genome BC (264PRO).

